# Cryo-EM structure of full-length HIV-1 Env bound with the Fab of antibody PG16

**DOI:** 10.1101/730333

**Authors:** Junhua Pan, Hanqin Peng, Bing Chen, Stephen C. Harrison

**Affiliations:** Laboratory of Molecular Medicine, Boston, MA 02115 USA; Howard Hughes Medical Institute, Boston Children’s Hospital, Boston, MA 02115 USA; Blackfan Circle, Boston, MA 02115 USA

**Keywords:** cryo-EM, cleaved gp160, broadly neutralizing antibody, immunogen design

## Abstract

The HIV-1 envelope protein (Env) is the target of neutralizing antibodies and the template for vaccine immunogen design. The dynamic conformational equilibrium of trimeric Env influences its antigenicity and potential immunogenicity. Antibodies that bind at the trimer apex stabilize a “closed” conformation characteristic of the most difficult to neutralize isolates. A goal of vaccine development is therefore to mimic the closed conformation in a designed immunogen. A disulfide-stabilized, trimeric Env ectodomain -- the “SOSIP” construct -- has many of the relevant properties; it is also particularly suitable for structure determination. Some single-molecule studies have, however, suggested that the SOSIP trimer is not a good representation of Env on the surface of a virion or an infected cell. We isolated Env (fully cleaved to gp120 and gp41) from the surface of expressing cells using tagged, apex-binding Fab PG16 and determined the structure of the PG16-Env complex by cryo-EM to an overall resolution of 4.6 Å. Placing the only purification tag on the Fab ensured that the isolated Env was continuously stabilized in its closed, native conformation. The Env structure in this complex corresponds closely to the SOSIP structures determined by both x-ray crystallography and cryo-EM. Although the membrane-interacting elements are not resolved in our reconstruction, we can make inferences about the connection between ectodomain and membrane-proximal external region (MPER) by reference to the published cryo-tomography structure of an Env “spike” and the NMR structure of the MPER-transmembrane segment. We discuss these results in view of the conflicting interpretations in the literature.

## Introduction

The HIV-1 envelope protein (Env), a trimer of gp160 subunits activated by cleavage to (gp120-gp41)_3_, is the sole surface antigen of the virus [1, 2]. Its dynamics, critical for its functions as the viral attachment protein and membrane fusogen, influence Env antigenicity by exposing epitopes inaccessible in the most stable equilibrium conformation [3–7]. Conformational fluctuations likewise influence the immunogenicity, both of viral Env during infection or propagation in culture, and of recombinant Env vaccine candidates.

The high rate of replication of HIV-1 in infected individuals, even during the long period of clinical latency, and the continuing host immune response together lead to great diversity in the amino-acid sequences of Envs derived from patient isolates [8]. The still unmet challenge for vaccine design and development has therefore been to devise a variant or series of variant Envs that can reliably elicit antibodies directed at certain reasonably conserved epitopes (so-called “broadly neutralizing” antibodies or bnAbs [9]). The finding that removing the transmembrane segment (TM) and cytoplasmic tail (CT) alters antigenicity of soluble, trimeric ectodomain implies that to focus on the conserved epitopes [10, 11], a vaccine immunogen may need to be a full-length Env trimer or a soluble Env ectodomain trimer stabilized in an appropriate conformation.

Early work comparing inhibition of infection by soluble CD4 (sCD4) with affinities of sCD4 for recombinant gp120 suggested an equilibrium between low CD4 affinity (“closed”) and high CD4 affinity (“open”) conformations [12], and many later structural studies of trimeric Env reinforced this notion [5, 13–15]. In the CD4-stabilized conformation, the V1-V2 loop flips over and moves outward, exposing the V3 loop and allowing formation of a “bridging sheet” between the inner and outer gp120 domains; accompanying gp41 rearrangements include repositioning of the fusion peptide [5, 13–15]. Intermediate, “partially open”, conformations have also been described [16]. The position of the equilibrium among these conformational states depends on Env sequence, as indicated by the early observation that sCD4 bound more readily to lab-adapted strains than to patient isolates and shown more completely by a recent detailed analysis of Env antigenicity [7, 17, 18]. The equilibrium for readily neutralized (“tier 1”) Envs favors the open conformation; that for “tier 2” and “tier 3” Envs lies toward the closed conformation [6, 7, 19, 20]. Envs that yield stable, homogeneous, trimeric, recombinant ectodomain (gp140) fall into the latter group [7, 17, 18]. Studies using fluorescence resonance energy transfer (FRET) or DEER spectroscopy have probed the ranges of relevant conformational ensembles [6, 21, 22].

High-resolution structures of soluble, trimeric HIV-1 Env have come from stabilizing the protein by introduction of a disulfide bond between the gp120 and gp41 fragments, mutation of an isoleucine to proline in gp41, truncating the gp41 ectodomain about 20 residues short of the transmembrane segment, and altering the cleavage site between gp120 and gp41 to ensure complete processing [4, 23, 24]. The resulting species, designated SOSIP.664 (the extension indicating the C-terminal residues of the soluble construct), has most of the antigenic properties of the same parent protein species on the surface of virions [25], but both BG505 SOSIP.664 and stable recombinant gp140 from two different isolates bind less strongly to antibodies that recognize the apex of trimer (variable loops V1-V2) than one might expect from the neutralization potency of those antibodies [17, 25]. The gp140 proteins also show greater exposure of variable loop 3 (V3) epitopes. These results suggest greater mobility of V1-V2 and V3 on the recombinant ectodomain than on the full-length trimer on virions. Subsequent studies confirmed this difference by showing that for cell-surface expressed Env, deletion of the CT or mutation of residues in the TM, as suggested by features of the NMR-determined structure of a bicelle-embedded, trimeric TM, led to a modified antigenic profile resembling that of the recombinant ectodomain [10, 11].

Do the antigenic differences just described reflect a broader dynamic range of the mutated or truncated species, or do the recombinant ectodomains have altered “ground state” conformations? We expect that potently neutralizing antibodies such as PG16 [26, 27] that bind tightly at the trimer apex of virion-borne Env will stabilize the conformation most prevalent on infectious virions. Spectroscopic distance measurements bear out this expectation [22]. To visualize the Env-PG16 complex directly, we have determined by cryo-EM, at 6Å resolution, the structure of a fully processed, full-length, cell-surface expressed Env, isolated directly by binding with affinity tagged PG16 Fab before extracting the complex from the cell membranes. We used a previously characterized, very stable Env from a clade A virus, 92UG037.8, with no modifications from the wild-type sequence [10]. We show that the structure of the Env ectodomain is essentially the same as that of BG505 SOSIP.664, which derives from a clade A virus with a closely related amino-acid sequence (82% sequence identity). The PG16 heavy-chain complementarity determining region 3 (HCDR3) shifts (relative to the rest of the Fab) from its position in complex with a scaffolded model for the V1-V2 region [27], so that the HCDR3 can fit between glycans projecting from two different Env subunits. The membrane proximal external region (MPER) and TM are not well ordered, and the CT appears to be fully disordered, probably because of the absence of a lipid-bilayer membrane. Nonetheless, low-resolution density features allow us to place a model from NMR of an MPER-TM trimer [28] and to suggest the connection of each gp41 chain between its ectodomain and MPER.

## Results

### Preparation of PG16(Fab)-gp160 complex

We expressed full-length HIV-1 Env from isolate 92UG037.8 on the surface of 293T cells and the PG16 Fab as a secreted product. We introduced a tandem strep II tag and a 10× histidine tag at the C-termini of heavy and light chains, respectively, incubated the Env expressing cells with the tagged PG16 Fab and used the affinity tags to purify the complex after detergent solubilization (see Methods). Size-exclusion chromatography gave a monodisperse complex eluting at a volume consistent with the size expected for a detergent-micelle solubilized gp160 trimer with a bound Fab (Fig. S1A). We have observed similar results using PG9 and PGT145, both of which bind, like PG16, at the trimer apex [29–31]. SDS-PAGE showed that the PG16 pulled out only the fully cleaved gp160 (Fig. S1B). The apparent stoichiometry is consistent with previous characterizations showing one Fab per trimer.

### Cryo-EM structure determination

We recorded image stacks (“movies”) for the PG16-gp160 complex both in open holes and, to circumvent problems with strong preferential orientation, in holes covered with a thin layer of continuous carbon. Figs. S2 and S3 show the image analysis scheme that led to a map at overall resolution of about 6.2 Å. To achieve a reasonably isotropic angular distribution, we initially eliminated a large fraction of the threefold views from the particle stack (Fig. S2). We then added back many of the particles and used updated processing procedures in Relion-3 [32] to achieve an overall resolution of about 4.5 Å. The estimated local resolution varied across the structure -- better than 5 Å in substantial parts of the core ectodomain and in the variable module of the Fab but poorer in some peripheral regions and completely ill defined for the TM and CT components. (Fig. S4). The Methods section describes both acquisition and analysis of the EM images.

### Model building

The Env amino-acid sequence of 92UG037.8 is very similar to that of BG505, the clade A isolate used for many of the published SOSIP structures (Fig. S5). We therefore docked the coordinates of an x-ray structure of glycosylated BG505 SOSIP(**PDB ID: 5V7J**) [33] into density corresponding to ectodomain in the sharpened 6.25 Å resolution map. We independently docked the PG16 Fab coordinates, from its complex (**PDB ID: 4DQO**) with the scaffolded V1-V2 from isolate ZM109 [27]. In the description of the gp160 trimer, below, we refer to the gp120 and gp41 fragments as chains A, C and E and B, D, and F, respectively, where each of the A-B, C-D, and E-F pairs derives from a single gp160 polypeptide chain.

There was diffuse density for the micelle surrounding the MPER-TM, but no well defined features that could place the MPER or TM within it (Fig. 1). Local classification of that region, after subtraction from each particle of the contribution from the rest of the molecule, yielded several classes with a shaft of density suggesting a location for the MPER-TM module (Fig. S6: compare classes 1,3 and 6, which together contain nearly 50% of all the particles). The orientation of the shaft of density was different among those classes for which it was prominent, suggesting a potentially flexible connection between the ectodomain and the entire membrane anchor. None of the classes gave density that could be fit with a single MPER-TM model, however. The corresponding linkage in influenza virus hemagglutinin is also flexible, but in that case, classification yielded a reasonable map for at least two different orientations of the TM segments relative to the ectodomain [34]. We could nonetheless introduce the model of a bicelle-embedded MPER-TM structure (**PDB ID: 6E8W**), determined by NMR [28], by aligning its known axial register with respect to phospholipid headgroups with the headgroup density in Env structures derived from cryo-electron tomography and subtomogram averaging [13] (Fig. 2A). We chose an approximate azimuthal orientation based on connections emanating from the C-terminal helices of the BG505 SOSIP.664 model (Fig. 2B) The CT, which includes several membrane-associated, amphipathic helices [35], is completely disordered, as expected in the absence of a lipid bilayer.

**Figure 1.**
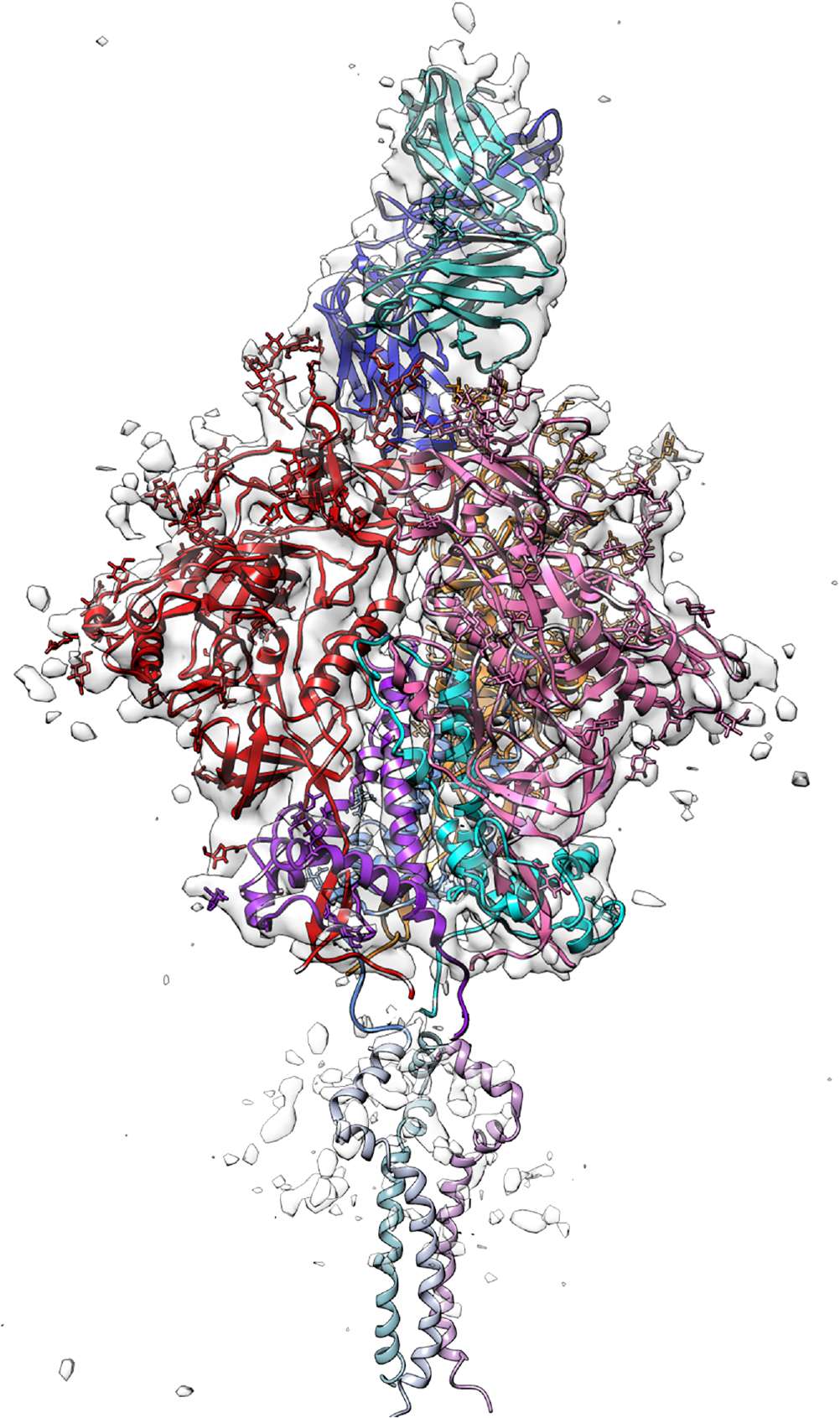
Cryo-EM reconstruction and model. Intermediate, 6.2 Å resolution reconstruction contoured to show outer profile of molecule, with model for ectodomain and PG16 Fab. Colors: gp120 in red (chain A), pink (chain C) and gold (chain E); gp41 in magenta (chain B), cyan (chain D) and blue (chain F); PG16 Fab in dark blue (heavy chain) and aquamarine (light chain). The MPER-TM model (**PDB ID: 6E8W**) [28], positioned as described in the text, is in fainter versions of the shades used to color the gp41 ectodomain (see also Fig. 3). Figure made with UCSF Chimera [49].

**Figure 2.**
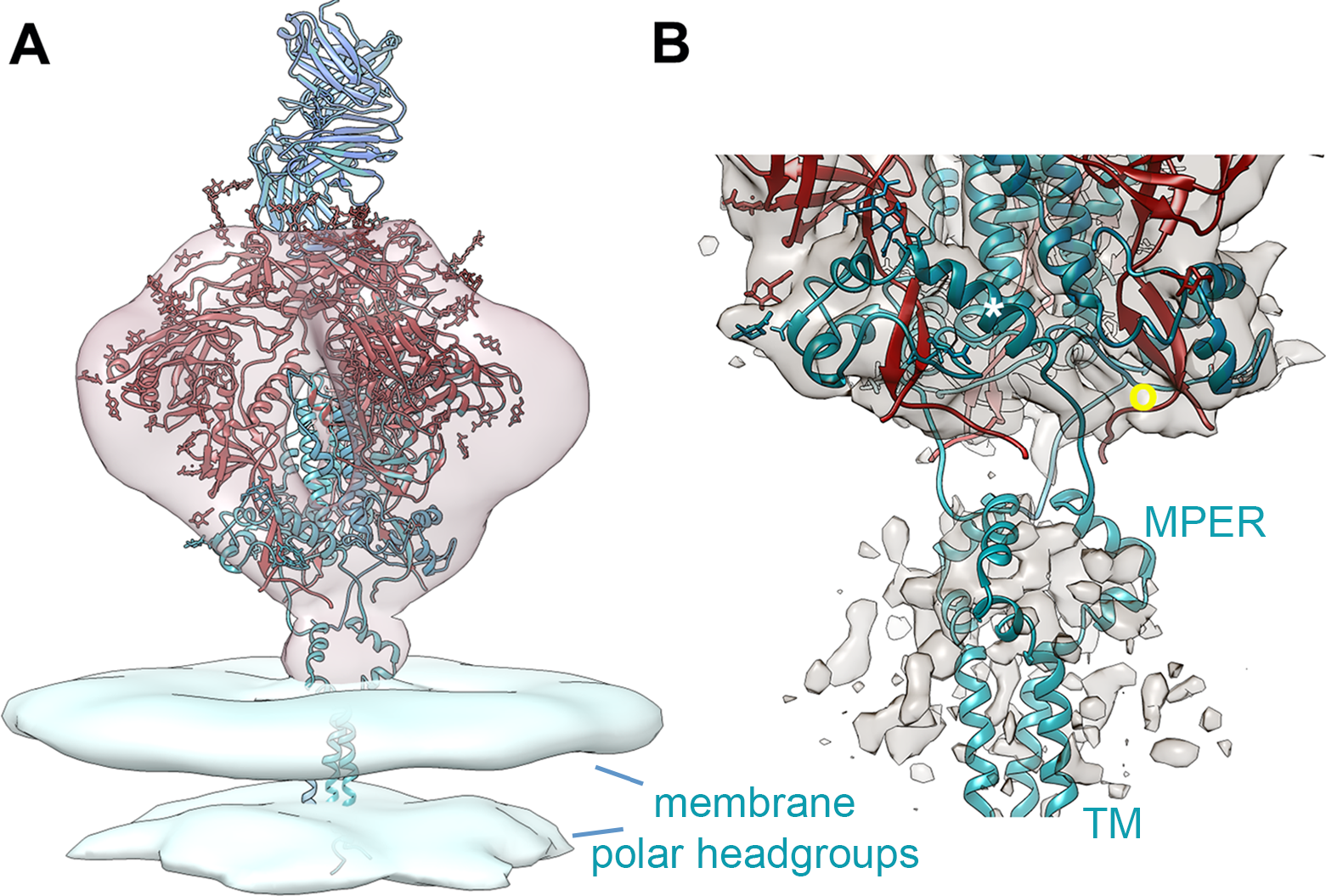
Comparison with low-resolution cryo-ET reconstruction of Env trimer on virion surface. (**A**) gp160 molecular model reported here fit into cryo-ET map (**EMDB ID: EMD5019**, pale red) [13], with MPER-TM model (**PDB ID: 6E8W**) [28] positioned in membrane density (**EMDB ID: EMD5022**, pale blue) [13] as described in Methods (see also Fig. 2D in reference [28]). Note that the relative positions of the two cryo-ET maps are fixed by the reconstruction, even though they are separate depositions. (**B**) Detail of the present reconstruction showing that helix 10 (white asterisk) does not continue directly into helix 11, as in the SOSIP structure, and that residues in SOSIP helix 11 instead connect into MPER. Yellow circle shows approximate position of SOSIP disulfide between chains C and D; helix 10 in foreground is on chain A. Density for MPER-TM region is dominated by contributions of detergent micelle, averaging of various orientations, and noise (Figs. S3 and S4); the azimuthal orientation of MPER-TM model around the threefold axis is therefore not well determined (see Methods). Figure made with UCSF Chimera [49].

### PG16 interactions

PG16 interacts with gp120 subunits A and E (Fig. 3). The density outline for HCDR3 shows that it retains, from its structure bound with a scaffolded V1-V2 (**PDB ID: 4DQO**) [27], an augmented β-sheet interaction between residues 100E to 100H and 167 to 171 of V2 on molecule A (Fig. 3A). The current resolution does not allow us to detect side-chain specificity, but the amino-acid sequence in this region of the isolate we used (92UG037.8) is similar to that of ZM109 (the isolate on which the scaffolded complex was based). In particular, conservation of charge on residues that participate in polar interactions (e.g., residues 168 and 171) is consistent with our inference from density that our PG16-gp160 complex and the PG16-scaffolded V1-V2 complex have the same hydrogen-bonding register. To avoid overlap elsewhere and to accommodate contacts from glycans at positions 160 on molecules A and E, at positions 134 and 156 on molecule A, and at position 188 on molecule E, the projecting HCDR3 loop (residues 99-100N), swings around by about 20° from its orientation with respect to the rest of V_H_ in the scaffolded variable region complex (Fig. 3B). The HCDR3 loop is disordered in crystals of uncomplexed PG9 and PG16 Fabs [29, 36], consistent with considerable flexibility.

**Figure 3.**
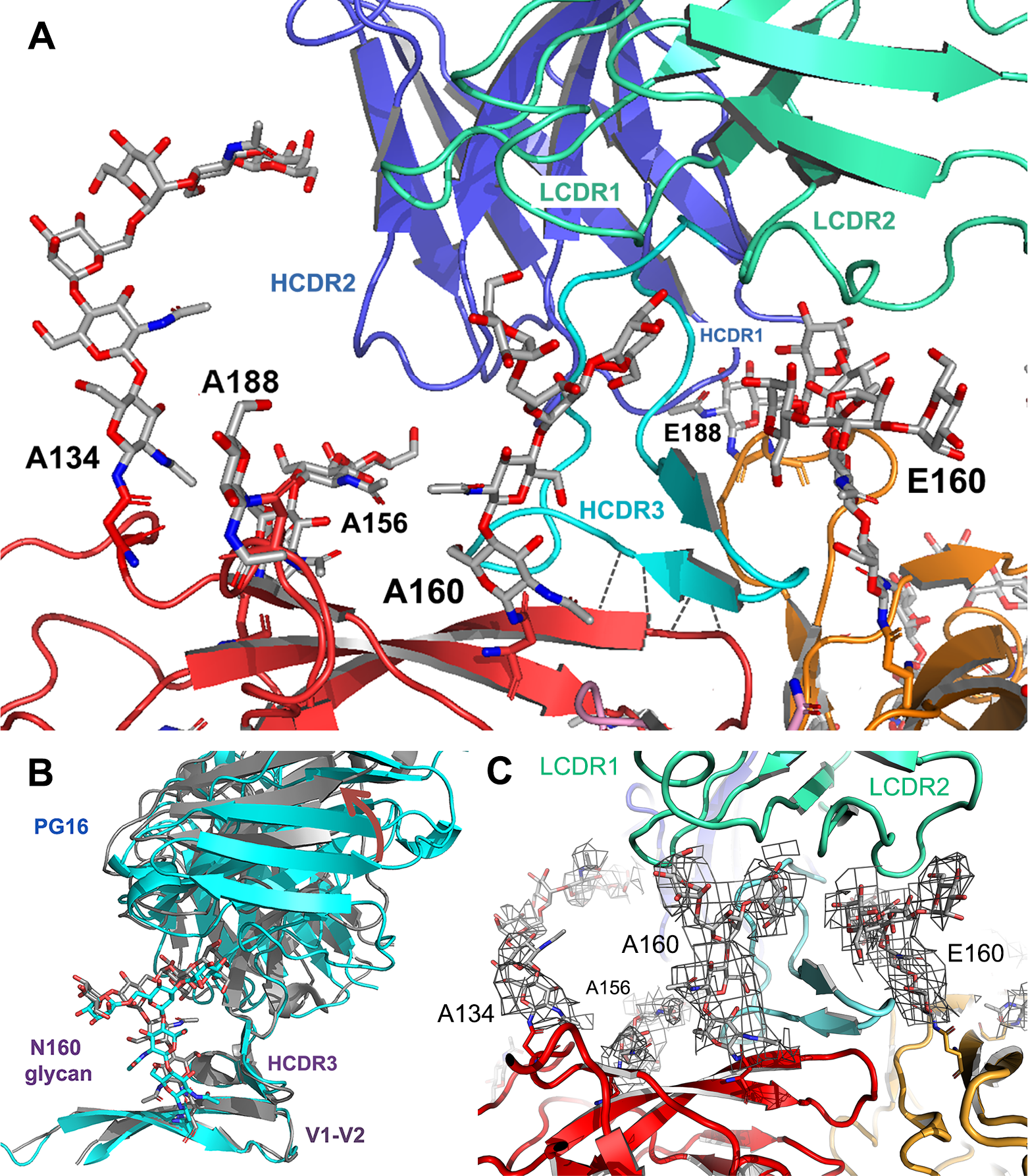
PG16 interaction. (**A**) Contact of PG16 Fab with apex of Env trimer. Viewpoint as in Fig. 1, but zoomed in so that chain C nearly vanishes in the foreground. Polypeptide chains are in ribbon representation; six glycans (labeled) are in stick representation. Colors as in Fig. 1, but with HCDR3 accented in cyan. (**B**) Superposition of PG16 and BC hairpin of V1-V2 in the gp160 structure (cyan) and scaffolded V1-V2 complex (**PDB ID: 4DQO**) [24] (gray), showing difference in orientation of the framework of the Fab variable module (brown arrow) with respect to the heavy-chain CDR3. The two structures are aligned on the V1-V2 BC loop. (**C**) As in (**A**), with contours for four glycans from final reconstruction at about 4.6Å resolution. Figures made with Pymol (Schrödinger, LLC).

Glycan contacts appear to fix the relative orientation of HCDR3 and the remainder of the variable module (Fig. 3). The glycan at Asn160 on chain A passes across the surface of PG16 HCDR3 to contact the light chain surface at LCDR1 and LCDR2 (Figs. 3A and 3C). The corresponding glycan on chain E passes across the other surface of HCDR3 to contact LCDR2. The glycan at Asn156 on chain A contacts HCDR2 (Figs. 3A and 3C). The outlines of the Asn160 density features, especially for the chain E glycan at 160 (Fig. 3C), are consistent with reported preference for mannose-rich glycans [27].

Two other glycans also appear to help position the PG16 variable-domain framework. One fills the prominent density feature extending from A-subunit Asn134 to the C” edge of the heavy-chain variable domain; the other bridges between the E-subunit Asn188 and the heavy-chain BC loop, with HCDR1 at its tip (Fig. 3A). The Asn 188 loop varies in length among different isolates; it is disordered in the BG505 SOSIP structure we used as a reference. We have built a rough model of the relatively short loop in our structure, but we cannot from our map make inferences about particular contacts.

Because most of the contacts between gp160 and PG16 other than those with HCDR3 appear to be glycan-mediated, it is possible that the “hinge” at the base of the HCDR3 loop that allows it to shift with respect to the variable-domain framework also allows the antibody to adapt to the alternative glycosylation patterns present on other isolates and to heterogeneity of glycan processing.

### Link with MPER

The C-terminal residue of the SOSIP model used as a starting point is at position 664. The NMR model for the MPER and TM (**PDB ID: 6E8W**) includes residues 660-710 [28]. Density for a long helix in the SOSIP model that comprises residues 636-664 (helices 10 and 11: Fig. S5) deviates, in our map, at about position 654 and extends into a thin connection into the MPER (Figs. 1 and 2B). The connecting sequence is completely polar: E^654^KNEQE^659^, and the MPER model, positioned as described above, begins immediately at 660.

The helical extension from 654 to 664 in the SOSIP.664 structures may be a consequence of the truncation at 664 and the SOSIP disulfide. The main chain of Cys 501, paired with Cys 605 in the SOSIP disulfide linkage, is in van der Waals contact with Leu663 (of the clockwise-related gp41) -- just where the gp41 polypeptide chain in the structure reported here deviates from the SOSIP conformation to join the MPER (Fig. 2B). Anchoring residue 501 probably fixes the otherwise somewhat flexible gp120 C-terminus at about position 505, thereby stabilizing the helical extension in the various SOSIP.664 constructs.

### Comparison with two other Fab-gp160 complexes

We have compared our structure with those for complexes of a cleaved, B-clade gp160 with Fabs from antibodies PGT145, which binds at the trimer apex (one Fab per trimer, somewhat like PG16, but with an even longer HCDR3)(**PDB ID: 6NIJ**) [37], and PGT151, which binds over the fusion peptide of gp41 (but only two sites per trimer occupied)(**PDB ID: 5FUU**) [38]. The structures of gp120-gp41 “protomers” from all three superpose very closely, despite the sequence differences that accompany clade divergence (Fig. S7), and hence all three superpose well on the BG505 SOSIP.664 backbone. The B-clade trimers are somewhat more asymmetric than A-clade 92UG037.8; it is plausible that their protomers shift around more strongly in response to asymmetric binding of the Fabs.

## Discussion

The structure of the gp160 ectodomain, as stabilized by association with a PG16 Fab, is essentially identical -- with a few local exceptions such as the most variable segments of V1-V2 and the connection to the MPER -- to that of the BG505 SOSIP.664. The stabilizing mutations introduced into the SOSIP ectodomain do not directly constrain V1-V2 or V3, but they presumably have indirect effects through enhanced stability of other parts of the structure. We infer that the antigenic differences at the trimer apex between SOSIP forms of soluble, recombinant gp140s and full-length, processed gp160 are due primarily to the extent to which they visit somewhat more open conformations, rather than to a difference in the most stable, “ground state” conformation. This conclusion acquires importance in view of on-going debate about whether the SOSIP structures indeed represent the conformation of native Env on infectious virions [21, 22].

That debate derives initially from interpretation of single-molecule FRET (smFRET) experiments, most of which have measured energy transfer between a fluorophore attached to an insert at about residue 136 in V1 and a fluorophore attached at about residue 400 in V4 of the same subunit [6]. The distance between the two positions is about 55 Å in our structure and in the BG505 SOSIP.664 gp140, but the fluorophores are on peptide insertions that may influence local conformational order-disorder equilibria (particularly in V4, which is generally quite disordered anyway) and both sites are also embedded in neighboring glycans. Moreover, the acceptor fluorophore is attached through a coenzyme A “handle” with a potential contour length of over 20 Å. The smFRET data have been analyzed into three “states”, with low (state 1), intermediate (state 3) and high (state 2) FRET signals, respectively [6, 21, 39]. Munro et al [6] reported that when Env on virion was bound with PG16 (and certain other broadly neutralizing antibodies), the low-FRET, state 1 predominated, but the same group reported recently that soluble SOSIP.664 instead gave a much higher FRET signal than did Env on virions and suggested that the SOSIP.664 structures determined by cryo-EM and x-ray crystallography do not represent the predominant conformation on the virion [21].

Measurements by DEER spectroscopy of inter-residue distances on two SOSIP.664 gp140s (from clade A BG505 and clade B, B41), reported last year, lead to a different conclusion [22]. The distances, maxima of relatively narrow distributions, are consistent with the various cryo-EM and x-ray structures of the stabilized species. Binding (separately) of two broadly neutralizing antibodies, 3BNC-117 and VRC34, does not change the peak distance measurements but in both cases appears to tighten up the distribution. That is, the SOSIP conformation that binds most tightly to those antibodies is indistinguishable, at the level of multiple inter-subunit distance measurements, from the directly determined structures.

The structure reported here comes from intact, fully cleaved gp160 pre-bound with PG16 Fab before extraction from the cell membrane and purified through an affinity tag on the Fab, thus ensuring continuous occupancy during the preparation. Previous work has shown that this species, when expressed on the surface of a cell, has an antigenic profile indistinguishable from its neutralization profile for the same or similar Envs on virions[10]. Those studies have also shown that truncation of the CT leads to some antigenic changes at the apex of the trimer, but to smaller or negligible antigenic changes elsewhere. Since we expect that the continuous presence of PG16 will have locked the apex of the trimer in its virion-borne conformation and since we see at the trimer base a well-packed structure that corresponds closely to the surface determined by sub-tomogram averaging of virus particles, we believe that the structure we see is a good representation of the principal conformation on a cell or a virion. Moreover, as it corresponds closely to the various SOSIP structures, we conclude that those structures are also good representations of virion-borne and cell-surface expressed Envs.

One way to reconcile this conclusion with the published smFRET data is to point out that depending on the orientation of the tether through which the acceptor fluorophore is attached, its distance from the donor fluorophore can range over ~ 30-40 Å, enough to span the difference between high and low FRET configurations. Interaction of virions and SOSIP gp140s with the passivated surface used for smFRET total internal reflection microscopy could in principle favor different configurations of the fluorophore, since in the former case, the labeled Env trimer could be a some distance from the surface, whereas in the latter, it is in direct contact. One would need to deconvolute the effects of fluorophore displacement from those of conformational excursions in order to make more definitive structural inferences from the smFRET results.

The antigenic properties of SOSIP gp140s do show that they sample a broader ensemble of conformations than do the corresponding, cell-surface expressed Envs [25, 40]. There are four principal differences between SOSIP.664 gp140 and Env on virions, any of which could influence the conformational dynamics of the protein: the disulfide between residues 501 and 605, the mutation of I559 to proline, the introduction of an ectopic cleavage site to ensure complete furin cleavage, and the truncation at residue 664. Residue 559 and the N-terminal end of the poorly ordered fusion peptide are potentially close together, and the effects of those two mutations might even correlate. Their location in the structure does not, however, suggest a role in longer-range conformational variability.

A correlation between the effects of truncation and disulfide linkage, because of the contact between residues 501 and 663 in SOSIP.664 structures, could influence conformational dynamics of membrane-proximal parts of the ectodomain. DEER spectroscopy measurements indeed show a broader distribution of inter-subunit distances at the base of the SOSIP trimer than at the apex [22], consistent with some loss of constraints from deletion of the MPER, TM and CT segments. The broader distribution indicates an increased dynamic range, but with the centroid of the principal peak still close to the position expected from SOSIP coordinates. Introduction of the SOSIP modifications into BG505 Env on virions also appears to broaden the distribution of occupancies, with a shift toward higher FRET states [21], but those measurements do not by themselves distinguish particular regions of the molecule.

In summary, we conclude from the structure reported here, and from structures described by others [41] [38], that the SOSIP.664 structures provide good pictures of the Env ectodomain on virions. Any single set of coordinates is necessarily a snapshot of a conformational ensemble. The range of SOSIP.664 conformational excursions appears to be broader than that of intact Env, but both center on a quite similar overall structure. Local deviations -- either static or dynamic -- can, of course, have substantial effects on antigenicity or immunogenicity of particular epitopes. Thus, the conformational similarity does not necessarily imply that SOSIP.664 trimers will be good vaccine components, but it does mean they are good structural guides to understanding the properties of authentic Envs.

## Materials and Methods

### Preparation of PG16 Fab

We cloned into a pVRC-IRES-puro vector (modified from a vector kindly provided by John Mascola, VRC, NIH) a codon-optimized gene (Genescript Piscataway, NJ) encoding both chains of the PG16 Fab with a 10X His and a tandem Strep II tag on the light and heavy chains respectively (each separated from its C-terminus by a GGGSA linker, with a GGGSGGGSGGSSA linker between the tandem tags) and established a stable cell line of HEK 293T cells by standard procedures. In brief, we used 5 μg plasmid DNA and Lipofectamine 3000 (Life Technologies, Grand Island, NY) to transfect 5×10^6^ cells in 10 mL DMEM supplemented with 10% FBS. After overnight incubation, cells were detached with trypsin-EDTA (Life Technologies, Grand Island, NY), split by series dilution, and monitored for single colonies. Colonies detected after 2-3 weeks were picked, protein expression estimated by Western blot, and high-expression clones were scaled up, distributed into 1 mL aliquots, frozen at −80°C in DMEM, 50% FBS, 1 μg/mL puromycin, 10% DMSO, and transferred to a liquid nitrogen storage tank.

For large-scale protein production, we thawed one 1 mL aliquot of the HEK 293T cells, established as just described, in 60 mL DMEM, 10% FBS, 1 μg/mL puromycin and expanded the cells after adaptation in suspension to Expi-293 medium (Life Technologies, Grand Island, NY) supplemented with 1 μg/mL puromycin. Fab was affinity purified from the medium using a Strepactin superflow resin (Abcam, Cambridge, MA) under gravity flow, followed by a Superdex 200 (GE Healthcare Bio-sciences, Pittsburgh, PA) size exclusion step.

### Preparation of PG16 (Fab) - HIV-1 gp160 complex

A stable cell line expressing HIV-1 gp160 from isolate 92UG037.8 has been described previously [10]. For protein preparation, we expanded cells from this line in 5-10 L Expi-293 medium to a density of 5-12×10^6^ cells/mL (verifying at >90% of cells were viable), centrifuged the cells gently at 300x g for 5-10 minutes, discarded the medium, dispersed the cell pellet in 400 mL 25 mM Tris pH 7.5, 150 mM NaCl, 1% BSA, incubated on ice with 5-10 mg PG16 Fab for 30 min, centrifuged at 300x g as above, and set aside the supernatant for recovery of excess PG16. We washed the Fab-bound cell pellet three times with 300 mL 25 mM Tris pH 7.5, 150 mM NaCl (buffer A), lysed the cells by incubating for 30 min in buffer A supplemented with 1% NP40, removed the cell debris by centrifugation at 10,000x g for 1 hr, and loaded the supernatant onto a 5 mL Strepactin superflow resin (Abcam, Cambridge, MA) run at 4°C under gravity flow. The resin was washed with 100 mL 25 mM Tris pH 7.5, 1 M NaCl, 0.025% NP40, then with 100 mL buffer A plus 0.025% NP40, before eluting with buffer A plus 0.025% NP40 and 2.5 mM desthiobiotin.

To reconstitute the PG16(Fab)-gp160 complex in Amphipol, we added to the pooled Strepactin elute a 10% solution of PMAL-C16 (Anatrace, Maumee, OH), dropwise with mixing, to a final PMAL-C16 concentration of 0.5%, and incubated on a rocker for at least 4 hr at 4°C. The sample was applied to a 1 mL Ni-NTA superflow column under gravity flow, the column washed twice with 2 L detergent-free buffer A, and the PG16(Fab)-gp160 complex eluted in 0.5 mL fractions in buffer A plus 300 mM imidazole. Fractions containing the complex were pooled, applied to a Superose 6 size-exclusion column (GE Healthcare Bio-sciences, Pittsburgh, PA), and eluted with buffer A (Fig. S1). When required, protein was concentrated on 50-100 μL NiNTA superflow resin and eluted with buffer A plus 500 mM imidazole in 100 μL fractions. Peak fractions were combined, and dialyzed in a dialysis button (Millipore, Burlington MA) overnight against 3 1L changes of buffer A to achieve a final concentration of ~1 mg/mL. The sample was diluted to 0.3 mg/mL for electron microscopy.

### Electron microscopy

We applied 3.2 μL PG16(Fab)-gp160 complex (at 0.3 mg/mL) to Quantifoil R 1.2/1.3 holey carbon 200-mesh copper grids (Quantifoil Micro Tools, Großlöbichau, Germany) or (at ~0.015 mg/mL) to C-flat R 1.2./1.3 holey carbon 400 mesh copper grids (Protochips, Morrisville, NC) coated with a thin layer of continuous carbon and plunge-froze in liquid ethane in a Vitrobot Mark 1 (FEI) with 4.2 sec blotting with filter paper pre-saturated at 100% humidity.

We recorded 2510 movies from the samples in open holes, with a K2 Summit detector and a Titan Krios electron microscopy (FEI) automated with SerialEM and operated at 300 kV and nominal magnification of 22,500x (calibrated pixel size 1.31 Å). Those images showed very strong preferential orientation along the gp160 threefold axis, and we therefore also recorded 5628 movies on a K2 Summit detector (Gatan, Inc., Pleasanton, CA) from samples on continuous carbon grids, using a Tecnai Polara electron microscope (FEI) automated with SerialEM [42] and operated at 300 kV and nominal magnification of 23,000x (calibrated pixel size 1.64 Å). In all cases, we recorded 40 frames/movie in super-resolution mode at a dose rate of 8 electrons/pixel/second and a total exposure of 40 electrons/Å^2^ (defocus range 1.2-3.0 μm).

### Image processing

We aligned movie frames with Unblur [43] and discarded those from thick ice or thick carbon (by inspection of aligned images 4x binned and low-pass filtered to 15 Å). Selecting 1000 aligned movies from the continuous carbon grids, we manually picked 287,298 particles with e2boxer [44], downsampled to 3.28 Å/pixel (with resample_mp in FREALIGN) [45], determined defocus with CTFFIND4 [46], and calculated 2D classes with RELION-2 [47] [48]. We discarded movies from poor 2D classes, leaving 202,859 particles for initial model generation with e2initialmodel[44]. We selected the best starting model using projection matching in IMAGIC and obtained alignments in RELION-2, followed by 10 cycles of Mode 1 refinement in FREALIGN [45] and several cycles of Mode 2. These procedures yielded a relatively isotropic, 8 Å resolution map with good definition of secondary structure. We used this map as the reference for later 3D alignments.

We used projections of this map (using e2projec3d from EMAN2), low-pass filtered to 20 Å resolution, as input for automated particle picking with gautomatch [K. Zhang, MRC LMB (www.mrc-lmb.cam.ac.uk/kzhang/)] from all remaining micrographs. We discarded particles with cross-correlation to the reference projection of less than 0.5, manually added apparently missed side views, and retained only particles from movies with Thon rings extending beyond 7 Å; the resulting data set contained 608,821 particles. We used resample_mp.exe (in Fourier space) [45] to resample the Krios images at a pixel size of 1.64 Å for merging with the more isotropically oriented particles from the Polara images (i.e., boxed the Krios particles to 320×320 pixels and downsampled to 256×256 pixels).

Because preferential views from the Krios data set dominated, we applied the procedure shown in Fig. S2 to smooth the distribution, ultimately extracting a total of 271,357 particles; 3D refinement of particles in this subset produced an apparently isotropic map at 6.4 Å resolution. The RELION classification scheme shown in Fig. S3 (left branch) yielded a final set of 125,075 particles and a resolution of 6.25 Å (Fig. S4). Sharpening with a B-factor of −300 Å^2^ produced the reconstruction used for initial interpretation and shown in Fig. 1. We then used the most recent version of Relion (v3.0.4) [32] for autorefinement of all 608,821 in the larger stack. It appeared to handle preferential orientation more robustly and yielded a reasonably isotropic map at 5.6 Å resolution. Classification into 6 classes produced a dominant (81%) class containing 492,995 particles with an overall resolution of 4.6 Å (Fig. S4). Further classification without alignment followed by autorefinement of the best class did not extend resolution further.

### Model building and refinement

We fitted the coordinates of BG505.SOSIP.664 (**PDB ID: 5V7J**) [8], PG16 Fab (**PDB ID: 4DQO**)[9] into the 6.25 Å resolution map with Chimera [49]. We used the sequence alignment (Fig. S5) for isolates BG505 and 92UG037.8 to generate 5000 structures using the Rosetta CM protocol [50], with secondary structure and disulfide restraints generated in PHENIX [51] and converted to Rosetta formats, selected the top 20 structures based on density scoring, and chose the best match to the map and reference model by visual examination, as a starting point for manual adjustment and refinement. Well-defined density for projecting glycans provided clear markers for the positions of modified asparagines. We built using O [52] a total of 87 glycans in the gp160 trimer and one in the PG16 light chain, paying attention to stereochemistry of published models [27, 33, 53] as well as bond lengths and bond angles [54]. Although lower contour levels often provided unambiguous evidence for at least three sugars (NAG-NAG-BMA) and in some cases for branching, we chose a relatively stringent cutoff, with most glycans built as just one or two sugars (NAG or NAG-NAG), and in a few cases, three. In the case of glycans that contact PG16, we could place branched structures even at high density cutoff, presumably because of order imposed by interaction with the Fab.

We carried out rigid body refinement and atomic displacement parameter (ADP) refinement in PHENIX [51], applying Ramachandran, rotamer, and secondary-structure restraints throughout, as well as non-crystallographic symmetry torsion-angle restraints for gp160. We then rebuilt glycans that had been incompletely constrained, to restore proper stereochemistry and glycosidic bond distances. These steps were then repeated for the 4.6 Å map; the polypeptide backbones did not shift significantly when fitting and refining the higher-resolution reconstruction.

### MPER and TM density

We analyzed density for the micelle-associated MPER and TM by creating masks that included only these regions. We subtracted signals beyond the mask boundary (using relion-project) and classified into 10 classes without further alignment, applying the 3D masks throughout. The classes, consisting of 13.37, 8.46, 15.76, 7.03, 8.82, 12.81, 7.33, 7.81, 9.76 and 8.86% particles, respectively, are shown in Fig. S6.

### Figure Preparation

We prepared the figures and videos using PyMOL (Schrödinger LLC), Chimera v1.13.1 [49], matplotlib and gnuplot. Multiple sequence alignment was performed with TCOFFEE [55] and displayed with ESPript [56].

### Accession numbers for atomic coordinates and maps

The 6.2 Å resolution cryo-EM map has been deposited at the EM Data Bank with EMDB ID EMD-20511, and the corresponding coordinates for the PG16-92UG037.8 Env complex have been deposited at the Protein Data Bank with PDB ID 6PWU. The 4.6 Å resolution map has been deposited at the EM Data Bank with EMDB ID EMD-20813, and the coordinates refined against that map have been deposited at the Protein Data Bank with PDB ID 6ULC.

## Acknowledgments

We thank Simon Jenni for advice on EM image processing, Zongli Li for advice and assistance with data collection on the Tecnai Polara at Harvard Medical School, and Chuan Hong and Zhiheng Yu at the HHMI EM facility for assistance with data collection on the Titan Krios at the Janelia Research Campus. The research was supported by National Institute of Allergy and Infectious Diseases Grant AI 100645 (Center for HIV/AIDS Vaccine Immunology -- Immunogen Discovery). S.C.H. is an Investigator in the Howard Hughes Medical Institute.

## References

[1] Wyatt R, Sodroski J. The HIV-1 envelope glycoproteins: fusogens, antigens, and immunogens. Science. 1998;280:1884–8.

[2] Harrison SC. Viral membrane fusion. Virology. 2015;479-480:498–507.

[3] Weissenhorn W, Dessen A, Harrison SC, Skehel JJ, Wiley DC. Atomic structure of the ectodomain from HIV-1 gp41. Nature. 1997;387:426–30.

[4] Pancera M, Zhou T, Druz A, Georgiev IS, Soto C, Gorman J, et al. Structure and immune recognition of trimeric pre-fusion HIV-1 Env. Nature. 2014;514:455–61.

[5] Ozorowski G, Pallesen J, de Val N, Lyumkis D, Cottrell CA, Torres JL, et al. Open and closed structures reveal allostery and pliability in the HIV-1 envelope spike. Nature. 2017;547:360–3.

[6] Munro JB, Gorman J, Ma X, Zhou Z, Arthos J, Burton DR, et al. Conformational dynamics of single HIV-1 envelope trimers on the surface of native virions. Science. 2014;346:759–63.

[7] Cai Y, Karaca-Griffin S, Chen J, Tian S, Fredette N, Linton CE, et al. Antigenicity-defined conformations of an extremely neutralization-resistant HIV-1 envelope spike. Proc Natl Acad Sci U S A. 2017;114:4477–82.

[8] Hemelaar J. The origin and diversity of the HIV-1 pandemic. Trends Mol Med. 2012;18:182–92.

[9] Kwong PD, Mascola JR, Nabel GJ. Broadly neutralizing antibodies and the search for an HIV-1 vaccine: the end of the beginning. Nat Rev Immunol. 2013;13:693–701.

[10] Chen J, Kovacs JM, Peng H, Rits-Volloch S, Lu J, Park D, et al. HIV-1 ENVELOPE. Effect of the cytoplasmic domain on antigenic characteristics of HIV-1 envelope glycoprotein. Science. 2015;349:191–5.

[11] Dev J, Park D, Fu Q, Chen J, Ha HJ, Ghantous F, et al. Structural basis for membrane anchoring of HIV-1 envelope spike. Science. 2016;353:172–5.

[12] Moore JP, McKeating JA, Huang YX, Ashkenazi A, Ho DD. Virions of primary human immunodeficiency virus type 1 isolates resistant to soluble CD4 (sCD4) neutralization differ in sCD4 binding and glycoprotein gp120 retention from sCD4-sensitive isolates. J Virol. 1992;66:235–43.

[13] Liu J, Bartesaghi A, Borgnia MJ, Sapiro G, Subramaniam S. Molecular architecture of native HIV-1 gp120 trimers. Nature. 2008;455:109–13.

[14] Wang H, Cohen AA, Galimidi RP, Gristick HB, Jensen GJ, Bjorkman PJ. Cryo-EM structure of a CD4-bound open HIV-1 envelope trimer reveals structural rearrangements of the gp120 V1V2 loop. Proc Natl Acad Sci U S A. 2016;113:E7151–E8.

[15] Harris A, Borgnia MJ, Shi D, Bartesaghi A, He H, Pejchal R, et al. Trimeric HIV-1 glycoprotein gp140 immunogens and native HIV-1 envelope glycoproteins display the same closed and open quaternary molecular architectures. Proc Natl Acad Sci U S A. 2011;108:11440–5.

[16] Wang H, Barnes CO, Yang Z, Nussenzweig MC, Bjorkman PJ. Partially Open HIV-1 Envelope Structures Exhibit Conformational Changes Relevant for Coreceptor Binding and Fusion. Cell Host Microbe. 2018;24:579–92 e4.

[17] Kovacs JM, Nkolola JP, Peng H, Cheung A, Perry J, Miller CA, et al. HIV-1 envelope trimer elicits more potent neutralizing antibody responses than monomeric gp120. Proc Natl Acad Sci U S A. 2012;109:12111–6.

[18] Kovacs JM, Noeldeke E, Ha HJ, Peng H, Rits-Volloch S, Harrison SC, et al. Stable, uncleaved HIV-1 envelope glycoprotein gp140 forms a tightly folded trimer with a native-like structure. Proc Natl Acad Sci U S A. 2014;111:18542–7.

[19] Montefiori DC, Roederer M, Morris L, Seaman MS. Neutralization tiers of HIV-1. Curr Opin HIV AIDS. 2018;13:128–36.

[20] Seaman MS, Janes H, Hawkins N, Grandpre LE, Devoy C, Giri A, et al. Tiered categorization of a diverse panel of HIV-1 Env pseudoviruses for assessment of neutralizing antibodies. J Virol. 2010;84:1439–52.

[21] Lu M, Ma X, Castillo-Menendez LR, Gorman J, Alsahafi N, Ermel U, et al. Associating HIV-1 envelope glycoprotein structures with states on the virus observed by smFRET. Nature. 2019;568:415–9.

[22] Stadtmueller BM, Bridges MD, Dam KM, Lerch MT, Huey-Tubman KE, Hubbell WL, et al. DEER Spectroscopy Measurements Reveal Multiple Conformations of HIV-1 SOSIP Envelopes that Show Similarities with Envelopes on Native Virions. Immunity. 2018;49:235–46 e4.

[23] Julien JP, Cupo A, Sok D, Stanfield RL, Lyumkis D, Deller MC, et al. Crystal structure of a soluble cleaved HIV-1 envelope trimer. Science. 2013;342:1477–83.

[24] Lyumkis D, Julien JP, de Val N, Cupo A, Potter CS, Klasse PJ, et al. Cryo-EM structure of a fully glycosylated soluble cleaved HIV-1 envelope trimer. Science. 2013;342:1484–90.

[25] Sanders RW, Derking R, Cupo A, Julien JP, Yasmeen A, de Val N, et al. A next-generation cleaved, soluble HIV-1 Env trimer, BG505 SOSIP.664 gp140, expresses multiple epitopes for broadly neutralizing but not non-neutralizing antibodies. PLoS Pathog. 2013;9:e1003618.

[26] Walker LM, Phogat SK, Chan-Hui PY, Wagner D, Phung P, Goss JL, et al. Broad and potent neutralizing antibodies from an African donor reveal a new HIV-1 vaccine target. Science. 2009;326:285–9.

[27] Pancera M, Shahzad-Ul-Hussan S, Doria-Rose NA, McLellan JS, Bailer RT, Dai K, et al. Structural basis for diverse N-glycan recognition by HIV-1-neutralizing V1-V2-directed antibody PG16. Nat Struct Mol Biol. 2013;20:804–13.

[28] Fu Q, Shaik MM, Cai Y, Ghantous F, Piai A, Peng H, et al. Structure of the membrane proximal external region of HIV-1 envelope glycoprotein. Proc Natl Acad Sci U S A. 2018;115:E8892–E9.

[29] McLellan JS, Pancera M, Carrico C, Gorman J, Julien JP, Khayat R, et al. Structure of HIV-1 gp120 V1/V2 domain with broadly neutralizing antibody PG9. Nature. 2011;480:336–43.

[30] Julien JP, Lee JH, Cupo A, Murin CD, Derking R, Hoffenberg S, et al. Asymmetric recognition of the HIV-1 trimer by broadly neutralizing antibody PG9. Proc Natl Acad Sci U S A. 2013;110:4351–6.

[31] Lee JH, Andrabi R, Su CY, Yasmeen A, Julien JP, Kong L, et al. A Broadly Neutralizing Antibody Targets the Dynamic HIV Envelope Trimer Apex via a Long, Rigidified, and Anionic beta-Hairpin Structure. Immunity. 2017;46:690–702.

[32] Zivanov J, Nakane T, Forsberg BO, Kimanius D, Hagen WJ, Lindahl E, et al. New tools for automated high-resolution cryo-EM structure determination in RELION-3. Elife. 2018;7.

[33] Zhou T, Doria-Rose NA, Cheng C, Stewart-Jones GBE, Chuang GY, Chambers M, et al. Quantification of the Impact of the HIV-1-Glycan Shield on Antibody Elicitation. Cell Rep. 2017;19:719–32.

[34] Benton DJ, Nans A, Calder LJ, Turner J, Neu U, Lin YP, et al. Influenza hemagglutinin membrane anchor. Proc Natl Acad Sci U S A. 2018;115:10112–7.

[35] Murphy RE, Samal AB, Vlach J, Saad JS. Solution Structure and Membrane Interaction of the Cytoplasmic Tail of HIV-1 gp41 Protein. Structure. 2017;25:1708–18 e5.

[36] Pancera M, McLellan JS, Wu X, Zhu J, Changela A, Schmidt SD, et al. Crystal structure of PG16 and chimeric dissection with somatically related PG9: structure-function analysis of two quaternary-specific antibodies that effectively neutralize HIV-1. J Virol. 2010;84:8098–110.

[37] Torrents de la Pena A, Rantalainen K, Cottrell CA, Allen JD, van Gils MJ, Torres JL, et al. Similarities and differences between native HIV-1 envelope glycoprotein trimers and stabilized soluble trimer mimetics. PLoS Pathog. 2019;15:e1007920.

[38] Lee JH, Ozorowski G, Ward AB. Cryo-EM structure of a native, fully glycosylated, cleaved HIV-1 envelope trimer. Science. 2016;351:1043–8.

[39] Ma X, Lu M, Gorman J, Terry DS, Hong X, Zhou Z, et al. HIV-1 Env trimer opens through an asymmetric intermediate in which individual protomers adopt distinct conformations. Elife. 2018;7.

[40] Pugach P, Ozorowski G, Cupo A, Ringe R, Yasmeen A, de Val N, et al. A native-like SOSIP.664 trimer based on an HIV-1 subtype B env gene. J Virol. 2015;89:3380–95.

[41] Torrents de la Peña A, Rantalainen K, Cottrell CA, Allen JD, van Gils MJ, Torres JL, et al. Similarities and differences between native HIV-1 envelope glycoprotein trimers and stabilized soluble trimer mimetics. bioRxiv. 2019:500975.

[42] Mastronarde DN. Automated electron microscope tomography using robust prediction of specimen movements. J Struct Biol. 2005;152:36–51.

[43] Grant T, Grigorieff N. Measuring the optimal exposure for single particle cryo-EM using a 2.6 A reconstruction of rotavirus VP6. Elife. 2015;4:e06980.

[44] Tang G, Peng L, Baldwin PR, Mann DS, Jiang W, Rees I, et al. EMAN2: an extensible image processing suite for electron microscopy. J Struct Biol. 2007;157:38–46.

[45] Grigorieff N. Frealign: An Exploratory Tool for Single-Particle Cryo-EM. Methods Enzymol. 2016;579:191–226.

[46] Rohou A, Grigorieff N. CTFFIND4: Fast and accurate defocus estimation from electron micrographs. J Struct Biol. 2015;192:216–21.

[47] Scheres SH. RELION: implementation of a Bayesian approach to cryo-EM structure determination. J Struct Biol. 2012;180:519–30.

[48] Fernandez-Leiro R, Scheres SH. Unravelling biological macromolecules with cryo-electron microscopy. Nature. 2016;537:339–46.

[49] Pettersen EF, Goddard TD, Huang CC, Couch GS, Greenblatt DM, Meng EC, et al. UCSF Chimera--a visualization system for exploratory research and analysis. J Comput Chem. 2004;25:1605–12.

[50] Song Y, DiMaio F, Wang RY, Kim D, Miles C, Brunette T, et al. High-resolution comparative modeling with RosettaCM. Structure. 2013;21:1735–42.

[51] Afonine PV, Poon BK, Read RJ, Sobolev OV, Terwilliger TC, Urzhumtsev A, et al. Real-space refinement in PHENIX for cryo-EM and crystallography. Acta Crystallogr D Struct Biol. 2018;74:531–44.

[52] Jones TA, Zou JY, Cowan SW, Kjeldgaard M. Improved methods for building protein models in electron density maps and the location of errors in these models. Acta Crystallogr A. 1991;47 (Pt 2):110–9.

[53] Gristick HB, von Boehmer L, West AP, Jr., Schamber M, Gazumyan A, Golijanin J, et al. Natively glycosylated HIV-1 Env structure reveals new mode for antibody recognition of the CD4-binding site. Nat Struct Mol Biol. 2016;23:906–15.

[54] Imberty A, Perez S. Stereochemistry of the N-glycosylation sites in glycoproteins. Protein Eng. 1995;8:699–709.

[55] Di Tommaso P, Moretti S, Xenarios I, Orobitg M, Montanyola A, Chang JM, et al. T-Coffee: a web server for the multiple sequence alignment of protein and RNA sequences using structural information and homology extension. Nucleic Acids Res. 2011;39:W13–7.

[56] Robert X, Gouet P. Deciphering key features in protein structures with the new ENDscript server. Nucleic Acids Res. 2014;42:W320–4.

[57] Kucukelbir A, Sigworth FJ, Tagare HD. Quantifying the local resolution of cryo-EM density maps. Nat Methods. 2014;11:63–5.

