## Supplemental Table S1 and Figs S1-S7 for "Cryo-EM structure of full-length HIV-1 Env bound with the Fab of antibody PG16"

**Table S1. Cryo-EM data collection, refinement and validation statistics**

| | | |
|---|---|---|
| **Data collection and processing** | | |
| Microscope | Tecnai Polara (HMS) | Titan Krios (JRC) |
| Magnification (x) (nominal/calibrated) | 23,000/30,487.80 | 22,500/38,167.94 |
| Voltage (kV) | 300 | 300 |
| Electron exposure (e–/Å$^2$) | 48 | 48 |
| Defocus range (μm) | 1.2-3.5 | 1.2-3.0 |
| Calibrated pixel size (Å) | 1.64 | 1.31 |
| Symmetry imposed | C1 | C1 |
| Initial subset particle images (no.) | 634,039 | 337,503 |
| Final subset particle images (no.)[#] | 80,628/212,176 | 44,447/280,819 |
| Final particle images (no.)[#] | 125,075/492,995 | |
| Map resolution (Å) (FSC threshold 0.143)[#] | 6.2/4.6 | |
| Map resolution range (Å)[#] | ∞-6.2/∞-4.6 | |
| **Refinement**[*] | | |
| Model resolution (Å) (FSC threshold 0.5) | 4.6 | |
| Map CC (%) | 0.6634 | |
| Model resolution range (Å) | ∞-4.6 | |
| Map sharpening *B* factor (Å$^2$) | -150 | |
| Model composition | | |
| Non-hydrogen atoms | 35,722 | |
| Protein residues | 2,301 | |
| Glycan/sugar residues | 88/142 | |
| *B* factors (Å$^2$) (overall/protein/glycan) | 153.52/154.71/142.14 | |
| *B* factors range (Å$^2$) | 78.12 - 435.31 | |
| R.m.s. deviations | | |
| Bond lengths (Å) | 0.004 | |
| Bond angles (°) | 0.744 | |
| Chirality | 0.071 | |
| Planarity | 0.003 | |
| Dihedral (°) | 7.100 | |
| **Validation**[*] | | |
| MolProbity score | 1.46 | |
| Clashscore | 4.84 | |
| Ramachandran plot | | |
| Favored (%) | 98.03 | |
| Allowed (%) | 1.97 | |
| Outliers (%) | 0.00 | |
| Rotamer outliers (%) | 0.44 | |
| C-beta deviations (%) | 0.00 | |
| Peptide Planes | | |
| Cis-Proline (%) | 3.37 (3 out of 89) | |
| Cis-general (%) | 0.00 | |
| Twisted-proline (%) | 0.00 | |
| Twisted-general (%) | 0.23 | |

[*]: Relion 2.0 with dominant view flattening/Relion 3.0.4.

[*]: Refinement and validation were reported by phenix.real_space_refinement [ref].

**Supplementary Figures**

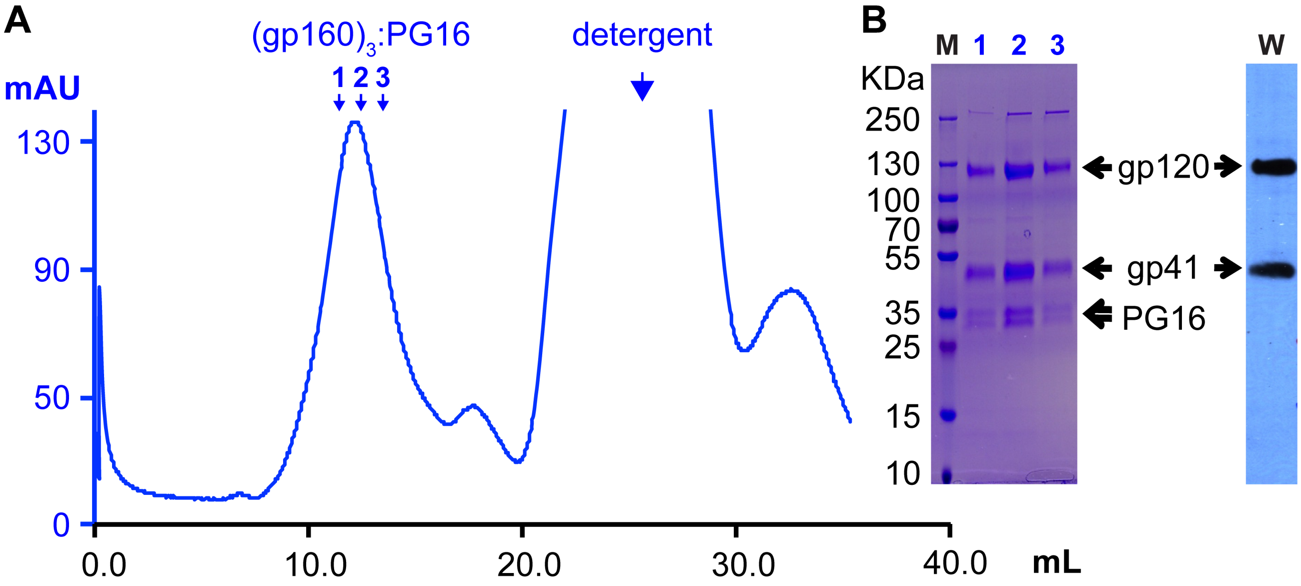


**Fig. S1**. Extraction of PG16-gp160 complex. (A) Size-exclusion chromatogram in buffer A (Superose 6 Increase 10/300 GL, bed volume 24 mL: GE Healthcare Bio-sciences, Pittsburgh, PA) after affinity elution from NiNTA (see Methods). (B) Left: Coomassie-stained gel of fractions indicated by arrows in the size-exclusion panel. Right: Western blot identifying gp120 and gp41. For the blot, electrotransfer to PVDF membrane at 100 V for 1 hr, in 25 mM Tris, 192 mM glycine, pH 8.3, 20% (v/v) methanol, was followed by 30 min incubation in blocking buffer (5% BSA in TBST) and 30 min incubation with antibodies 240D and Ab1281 (1 μg/mL in blocking buffer). The membrane was then washed three times for 5 min each in TBST, followed by a 30 minute incubation with horseradish peroxidase (HRP) conjugated goat anti-human secondary antibody (0.5 μg/mL in blocking buffer) and 3 washes with TBST. The blot was then soaked in 1 mL premixed Immun-Star HRP detection solution (Bio-Rad) for 1 min and exposed to film for 5 sec.

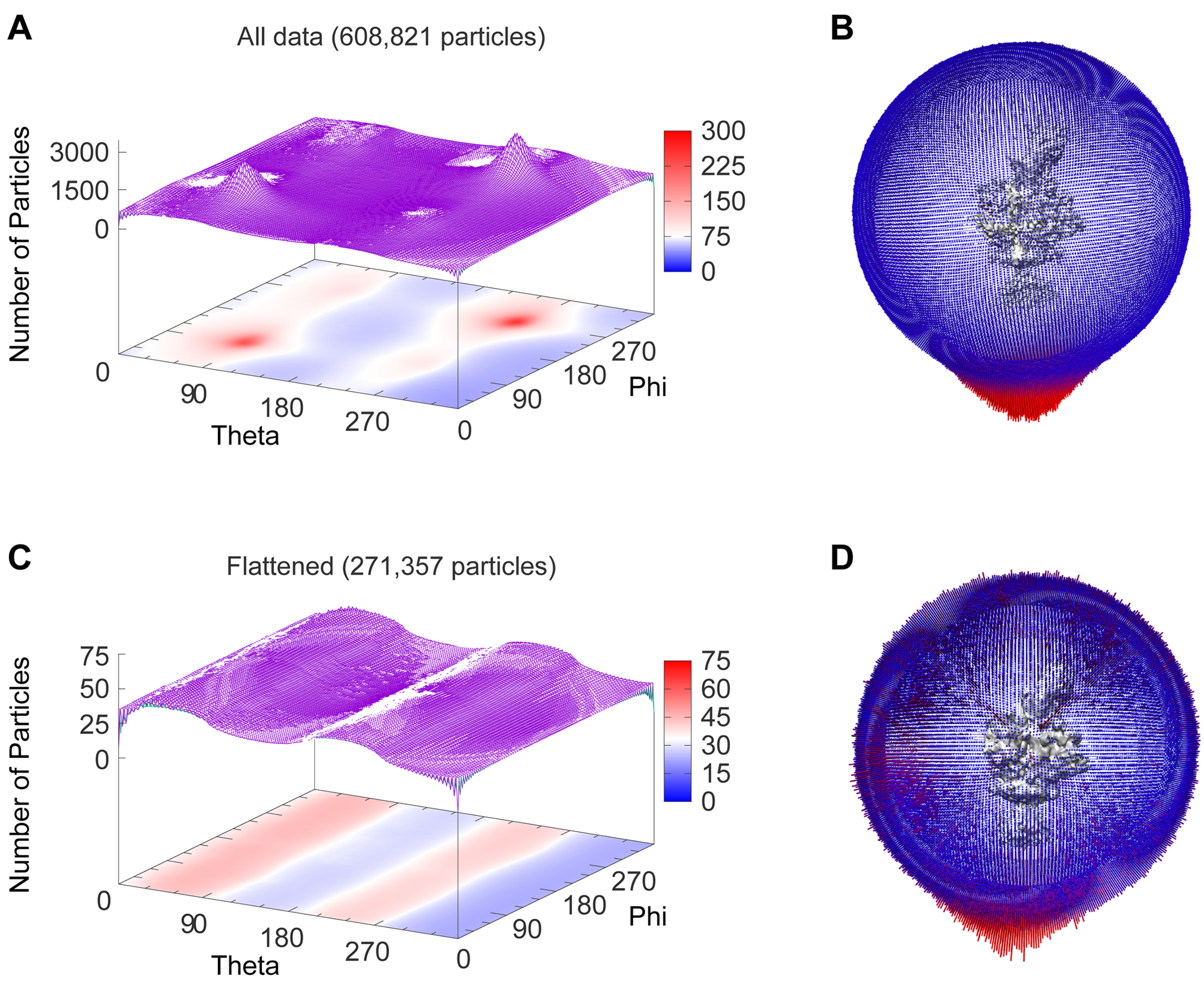


**Fig. S2**. Angular distribution analysis. (A) Plot of theta/phi distribution, shown as both a "landscape" and a "heat map". In the heat map, red was set as ≥300 particles for a theta-phi pixel. As shown in the landscape plot, the actual value for the preferred orientations was about 3000. There is a very strong preference for 90°/90° and 270°/270°, i.e., views along the threefold axis of gp160. (B) Corresponding plot from Relion [47]. (C) and (D) Distribution after "flattening". There is still some preference for views near the gp160 threefold. In the heat map, red represents 75 particles for a theta-phi pixel.

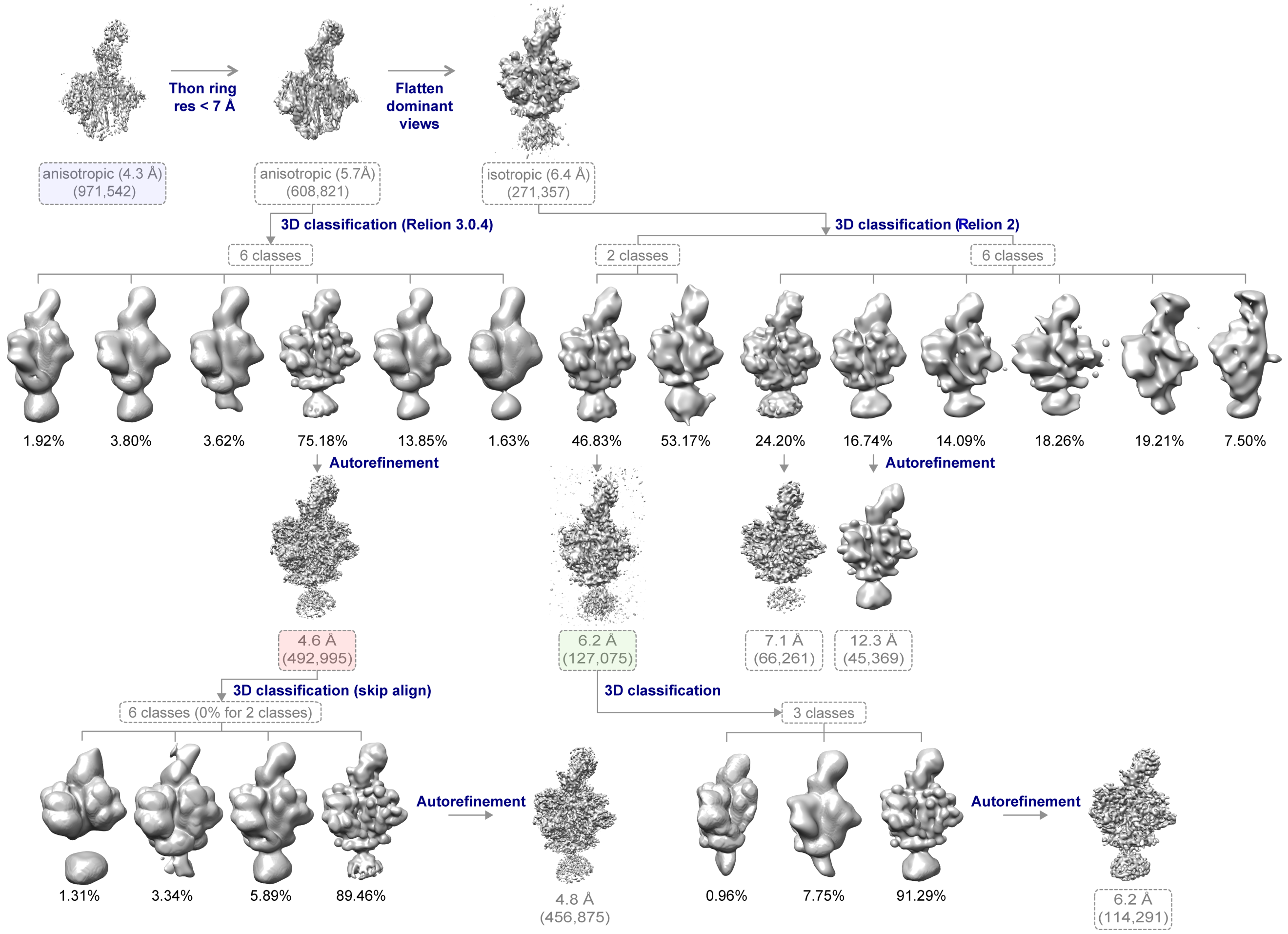


**Fig. S3**. 3D classification and refinement scheme. After eliminating frames with Thon rings that did not extend beyond 7 Å (which in practice corresponded largely to poor images and hence served more as a quality filter than as a resolution criterion) and flattening the angular distribution by removing a fraction of the dominant views (Fig. S2), we subjected 271,357 particles to classifications with 2 and 6 classes, respectively. Including the four additional classes did not improve the resolution of the best class, and we therefore selected the better of the classes from the two-class refinement, removed particles based on subclassification as shown, and carried out autorefinement on the remaining 114,291 particles. We used the resulting reconstruction, at 6.2 Å resolution, for initial model building. When Relion-3 became available, we returned to the particle stack after Thon-ring filtering and carried out the sequence of classifications and refinements shown. The contours in Fig. 3C are from the reconstruction at 4.6 Å resolution (red highlight). Further classification did not improve the resolution, as indicated by the steps at the bottom left.

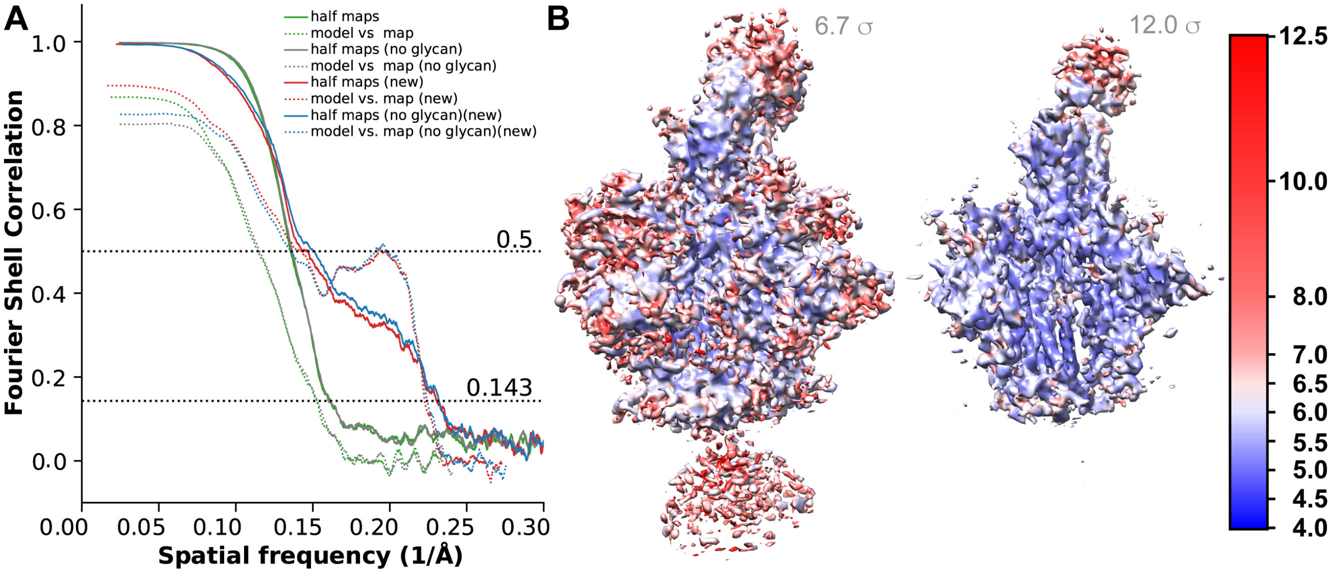


**Fig. S4**. Resolution analysis. (A) Fourier shell correlation analysis. We show the results for both the 6.2 Å reconstruction (green and gray curves) and the subsequent 4.6 Å reconstruction (red and blue curves: labeled "new" in panel). Half-map FSCs in solid lines; model-vs.-map FSCs in dotted lines. We show calculations for both the entire structure and for a volume lacking the glycan features. (B) Local resolution calculated with ResMap [57]. The two images show different contouring levels in Chimera [49].

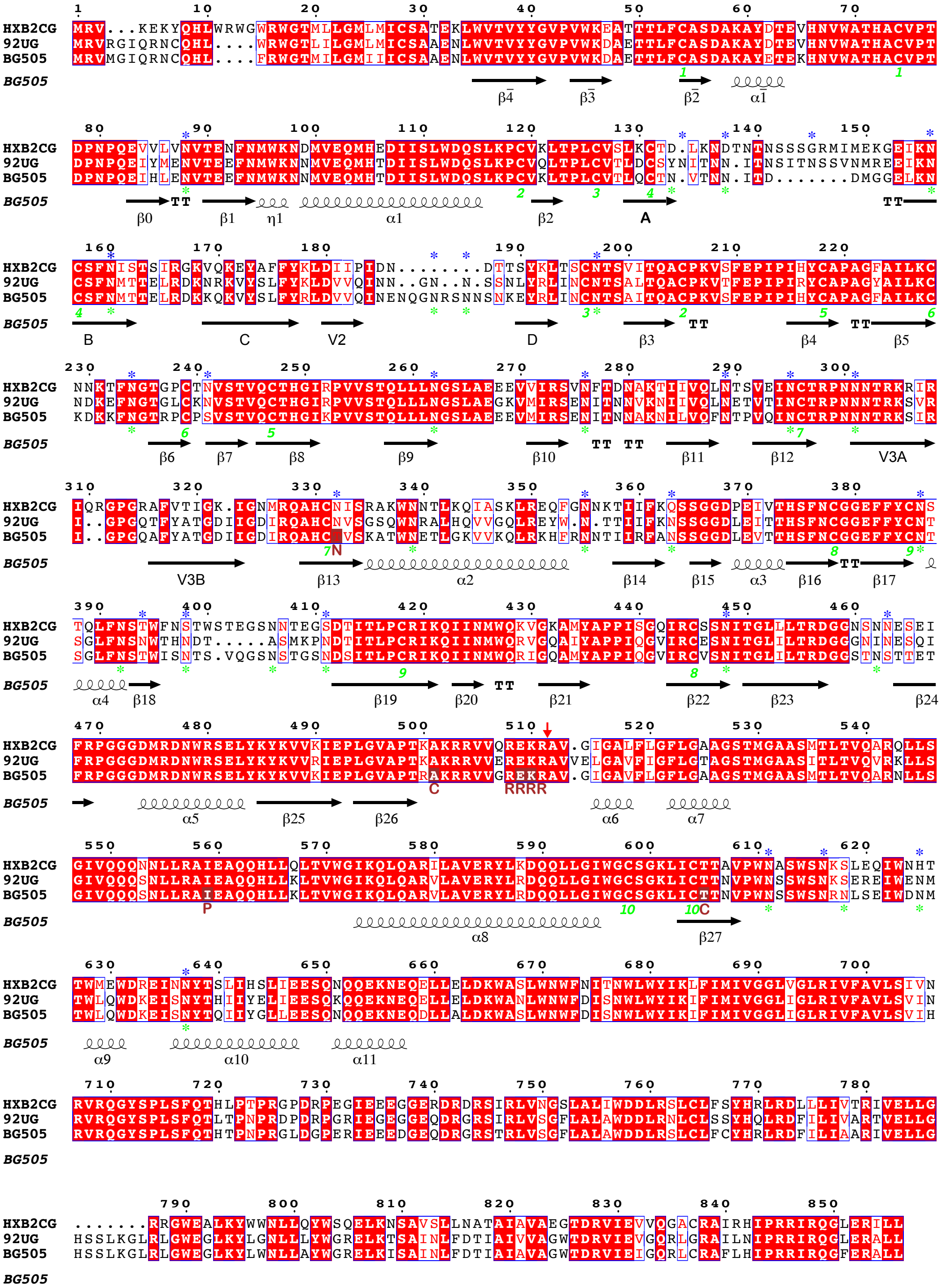

**Fig. S5**. Alignment of Env sequences: BG505 and 92UG037.8 aligned with HXB2 numbering standard. Conventional secondary structure designations are shown beneath the alignment. Blue asterisks show potential N-glycosylation sites for BG505; green asterisks, for 92UG037.8. Green numerals indicate disulfides between cysteines of the same number. Red highlights show conserved residues; red letters indicate physical-chemical conservation. Letters in dark red below the BG505 sequence show modifications in the SOSIP construct.

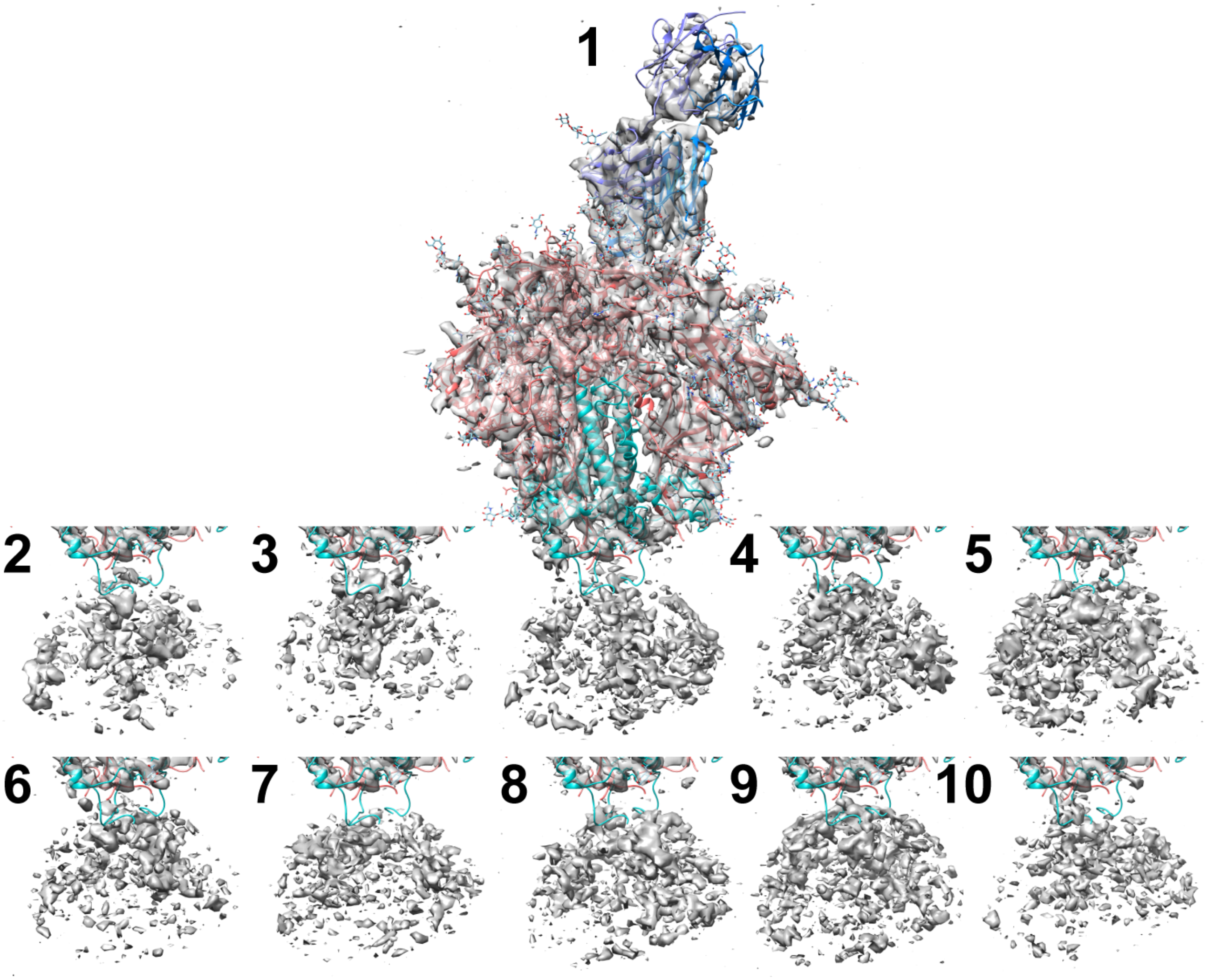


**Fig. S6**. Local classification, with signal subtraction, of the volume containing MPER, TM, and detergent micelle. Class 1 is in the center, aligned with the ecotodomain density; the other classes, contoured similarly, are shown alone. Classes 1, 3 and 6 (the three most populated) have central density potentially attributable to the MPER-TM, but in different orientations with respect to the ectodomain. The percentages of total particles in each class are: 13.37, 8.46, 15.76, 7.03, 8.82, 12.81, 7.33, 7.81, 9.76 and 8.86%, for classes 1 through 10, respectively.

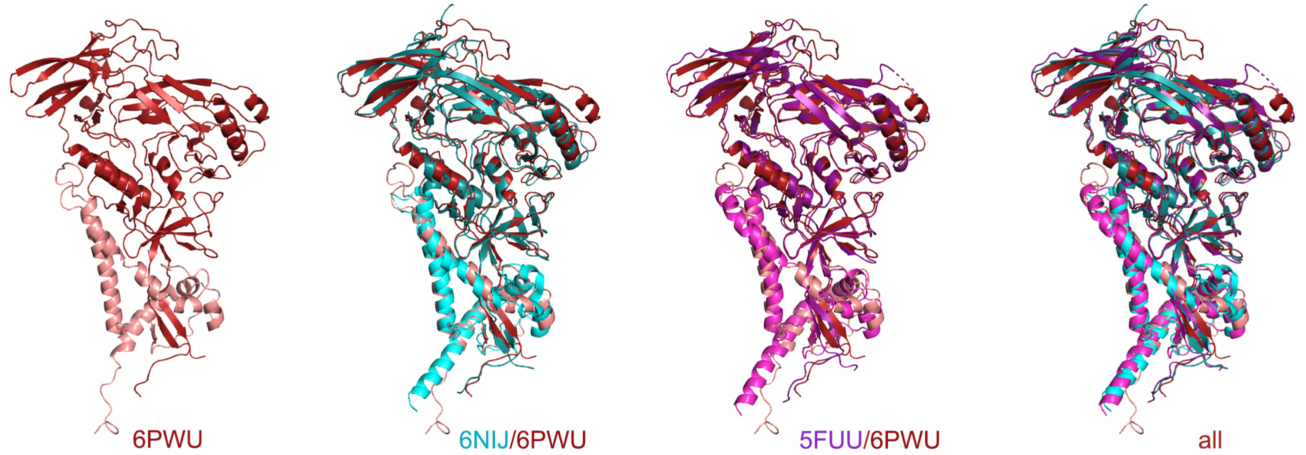


**Fig. S7**. Comparison of the structure reported here with those of cleaved gp160 bound with PGT145 (PDB:6NIJ) and cleaved gp160ΔCT bound with PGT151 (PDB:5FUU). The superpositions shown here were calculated using the core regions of gp120, for a single gp120/gp41 protomer (the A/B protomer in all three cases).
